# A Water Relaxation Atlas for Age- and Region-specific Metabolite Concentration Correction

**DOI:** 10.1101/2024.09.27.615424

**Authors:** Gizeaddis Lamesgin Simegn, Yulu Song, Saipavitra Murali-Manohar, Helge J. Zöllner, Christopher W. Davies-Jenkins, Dunja Simičić, Kathleen E. Hupfeld, Aaron T. Gudmundson, Emlyn Muska, Emily Carter, Steve C.N. Hui, Vivek Yedavalli, Georg Oeltzschner, Douglas C. Dean, Can Ceritoglu, Tilak Ratnanather, Eric Porges, Richard Edden

## Abstract

Metabolite concentration estimates from Magnetic Resonance Spectroscopy (MRS) are typically quantified using water referencing, correcting for relaxation-time differences between metabolites and water. One common approach is to correct the water reference signal for differential relaxation within three tissue compartments (gray matter, white matter, and cerebrospinal fluid) using fixed literature values. However, water relaxation times (*T*_1_ and *T*_2_) vary between brain locations, in pathology, and with age. MRS studies, even those measuring metabolite levels across the lifespan, often ignore these effects, because of a lack of reference data. The purpose of this study is to develop a water relaxometry atlas and to integrate location- and age-appropriate relaxation values into the MRS analysis workflow. 101 volunteers (51 men, 50 women; ∼10 male and 10 female participants per decade (from the 20s to 60s), were recruited. *T*_1_-weighted MPRAGE images ((1-mm)^3^ isotropic resolution) were acquired. Whole-brain water *T*_1_ and *T*_2_ measurements were made with DESPOT ((1.4 mm)^3^ isotropic resolution). *T*_1_ and *T*_2_ maps were registered to the JHU MNI-SS/EVE atlas using affine and LDDMM transformation. The atlas’s 268 parcels were reduced to 130 by combining homologous parcels. Mean *T*_1_ and *T*_2_ values were calculated for each parcel in each subject. Linear models of *T*_1_ and *T*_2_ as functions of age were computed, using *age – 30* as the predictor. Reference atlases of “age-30-intercept” and age-slope for *T*_1_ and *T*_2_ were generated. The atlas-based workflow was integrated into Osprey, which co-registers MRS voxels to the atlas and calculates location- and age-appropriate water relaxation parameters for quantification.

The water relaxation aging atlas revealed significant regional and tissue differences in water relaxation behavior across adulthood. Using location- and subject-appropriate reference values in the MRS analysis workflow removes a current methodological limitation and is expected to reduce quantification biases associated with water-referenced tissue correction, especially for studies of aging.

## 1. Introduction

The most common approach for quantifying metabolite concentrations in vivo with proton magnetic resonance spectroscopy (^1^H MRS) is water referencing, that is, assuming that the ratio between metabolite concentrations and tissue water concentration can be inferred from the ratio of metabolite signals to the unsuppressed water signal from the same tissue volume (1). This approach is the subject of much prior literature (2–5), engaging various degrees of sophistication to correct for known measurement biases. For example, it is known that metabolite and water signals have different relaxation times, both transverse (*T*_2_) and longitudinal (*T*_1_), therefore relaxation correction is commonly applied. It is known that water within different tissues has different *T* _1_s and *T*_2_s, so tissue effects are also often considered. Most modern MRS analysis packages (6–8) co-register the MRS voxel location to structural *T*_1_-weighted images to segment the volume of interest into gray matter (GM), white matter (WM), and cerebrospinal fluid (CSF). It is therefore possible to correct the acquired water reference signal for differential relaxation within these three compartments, assuming literature values for each.

The Gasparovic method (1) uses 11 literature values to infer metabolite concentration: the MR-visible water concentration (or visibility fraction) in GM, WM, and CSF (*c*_wi_ where the subscript *i* corresponds to each of the three tissue compartments); the *T*_1_ of water in GM, WM, and CSF (*T*_1wi_); the *T*_2_ of water in GM, WM, and CSF (*T*_2wi_); and the *T*_1_ and *T*_2_ of the metabolite signal (*T*_1met_ and *T*_2met_). These are combined with five measured parameters: the metabolite signal (or linear combination model component) *S*_met_, the water signal amplitudes *S*_w_, and the volume fraction of GM, WM, and CSF (f_vol,i_). In line with community consensus practice, toolboxes using water-scaled and tissue-corrected quantification such as Osprey software (6) take 3T reference values for water relaxation using several widely used references. Because segmentation is only typically considered at the level of GM, WM, and CSF, the same relaxation values would be applied for gray matter in the cortex and thalamus, for example, regardless of the known differences in relaxation across the brain within each tissue type.

Similarly, extensive literature shows changes in water relaxation with age. For example, it is a common observation that GM-WM contrast in *T*_1_-weighted structural images diminishes with age. Both longitudinal *T*_1_ (9–12) and transverse *T*_2_ (13–16) relaxation times change across the adult lifespan. Such changes are widely ignored in MRS studies, even MRS studies that explicitly seek to measure changes in metabolite levels across the lifespan. In the absence of appropriate correction, age-dependent relaxation changes spuriously present themselves as age-dependent changes in metabolite concentrations. This is a confounding factor for many MRS studies, particularly those conducted across the lifespan.

Therefore, the goal of this project is to address these limitations in quantification practice by acquiring a water relaxometry dataset in a structured cohort of subjects across the adult lifespan from 20–60 years of age. Non-linear co-registration to a structural atlas allows us to summarize this dataset at the level of 130 structural parcels and to test for changes in water *T*_1_ and *T*_2_ with age. Linear models in each anatomical parcel give age-30 intercept values of *T*_1_ and *T*_2_ and age-slopes, which have been summarized as a water relaxometry atlas and integrated into Osprey to allow future MRS analyses to use region-, and age-appropriate water relaxation reference values.

## 2. Methods

### 2.1. Cohort Description

101 volunteers were recruited and provided their written informed consent to participate. The Johns Hopkins University and University of Florida Institutional Review Boards approved all study procedures. The cohort was structured to include approximately ten male and ten female participants in each decade: 20s; 30s; 40s; 50s; and 60s. Half of the cohort (51 subjects) were recruited and scanned at Johns Hopkins, and the remainder were recruited and scanned at the University of Florida, Gainesville.

### 2.2. Acquisi’on Methods

*T*_1_-weighted MPRAGE images were acquired with 1 mm^3^ isotropic resolution, with scan TR 3.7 ms, TE 8.1 ms, and FA 8°. The DESPOT (driven-equilibrium single-pulse observation of *T*_1_/*T*_2_) method (17) was used for rapid whole-brain measurement of *T*_1_ and *T*_2_. *T*_1_ was measured from a series of spoiled gradient-recalled echo (SPGR) images acquired at flip angles of 4°, 12°, and 18°. This *T*_1_ information was combined with the fully balanced steady-state free precession (bSSFP) images acquired at flip angles of 15°,30°, and 60° at two phases cycling patterns (0°, 180°) to determine T_2_. DESPOT images were acquired with FOV 224×224×155 and (1.4 mm)^3^ resolution.

### 2.3. Analysis Methods

The acquired DESPOT images (SPGR and bSSFP), and the *B*_1_ map were linearly co-registered to correct for head movement (18), and non-parenchyma signals were excluded (19). The DESPOT1 and DESPOT2-FM techniques were applied for *T*_1_ and *T*_2_ mapping (20). *T*_1_ was estimated from SPGR data using flip angles, and *T*_2_ was derived from bSSFP images using a fitting approach based on different flip angles. NIFTI-format *T*_1_ and *T*_2_ maps were generated in a Python environment via the qmri-neuropipe tool(21). *T*_1_-weighted images were co-registered to the JHU atlas (i.e. JHU MNI-SS/EVE (22)) by computing an affine and LDDMM (23–25) transformation, as shown in Figure 1, and those transforms applied to the DESPOT *T*_1_ and *T*_2_ maps, resulting in standard-space quantitative *T*_1_ and *T*_2_ images.

**Figure 1.**
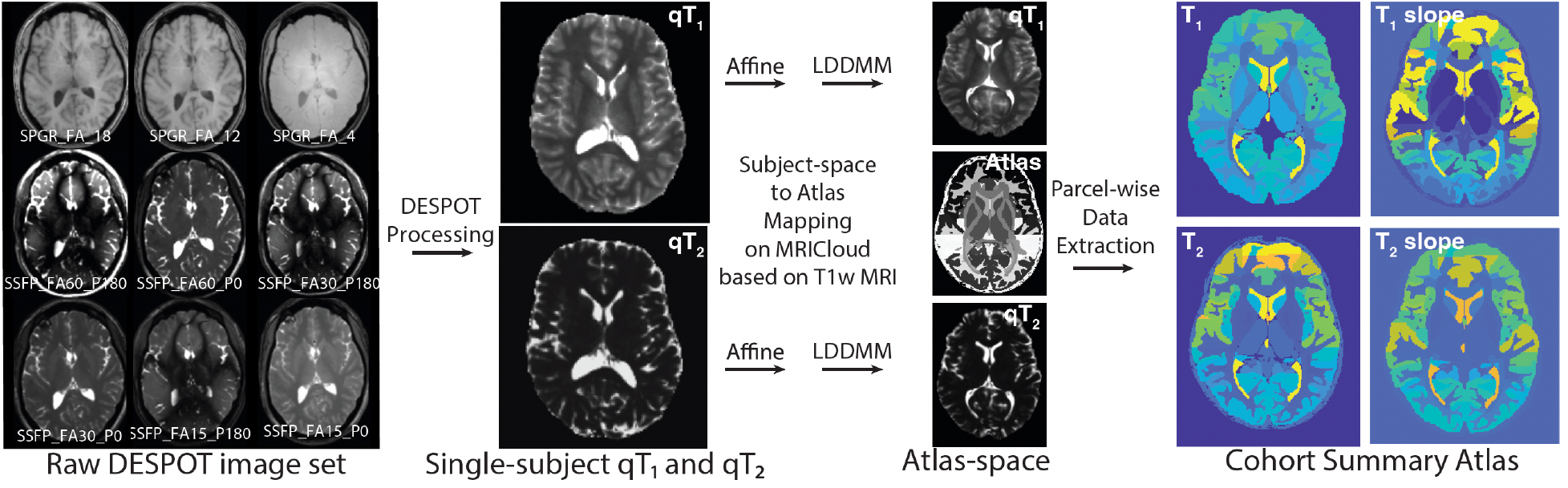
Flowchart illustrating the process of creating the relaxometry atlas.

### 2.4. Statistical Analysis

The 268 parcels in the JHU atlas were reduced by combining homologous left- and right-side parcels, and all internal CSF spaces into a single parcel, yielding 130 parcels. For each of the 130 parcels, mean *T*_1_ and *T*_2_ values were calculated for every subject. Linear models were then computed for *T*_1_ and *T*_2_ individually in each parcel, with (subject age – 30) as the predictor and two fitted parameters: slope and intercept. Using MATLAB 2024a, regression models were applied to fit the data, estimating both slope and intercept values. Additionally, p-values for the t-statistic of a two-sided test, assessing whether the slope was significantly different from zero, and R^2^ values for model fit goodness were reported. In order to generate usable intercepts that do not extrapolate beyond the dataset, age–30 was used as the predictor so that model intercepts would represent predicted relaxation times at 30 years of age (rather than 0 years of age) (16).

An age-30 reference atlas for *T*_1_ and *T*_2_ was then populated by filling each parcel in a standard-space atlas with the appropriate intercept value. An age-slope atlas was similarly populated, allowing predicted *T*_1_ and *T*_2_ values to be calculated for each parcel given a subject age. For example, the predicted *T*_1_ value for a given parcel, *j, T* _1j,predicted_ can be calculated as:

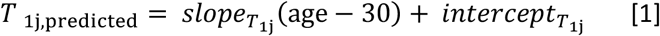

These values were also collected in a look-up table (supplementary Table S1).

### 2.5. Integration into Osprey

Within this dataset, we apply the non-linear LDDMM deformation to deliver atlas segmentation for each subject. However, this is a time-consuming cloud-based process that is not envisaged for typical future Osprey analyses. Therefore, we have developed a workflow to leverage the relaxometry atlas for future analyses of MRS datasets that do not have single-subject atlas segmentation. Resources for this workflow are stored in the source code directory osprey/quantify/atlas. The 130-parcel JHU atlas image of parcel labels was co-registered to SPM standard space using SPM ‘Estimate’, i.e. without re-slicing the discrete image, and storing the resulting image as ‘atlas_130.nii’. For clarity, we will here refer to the spectroscopy voxel as the voxel, and the individual elements of images as microvoxels.

While it is possible to generate relaxometry atlas images, such as those shown in Figure 2, the most efficient way to store these data in Osprey is as a single atlas image of parcel numbers and a look-up table, which is stored as ‘*AtlasLookUpTable*.*mat*’. For each numbered atlas parcel, this file includes anatomical labels, tissue allocations (GM or WM), and *T*_1_ and *T*_2_ intercepts and slopes (as in Supplemental Table S1).

**Figure 2.**
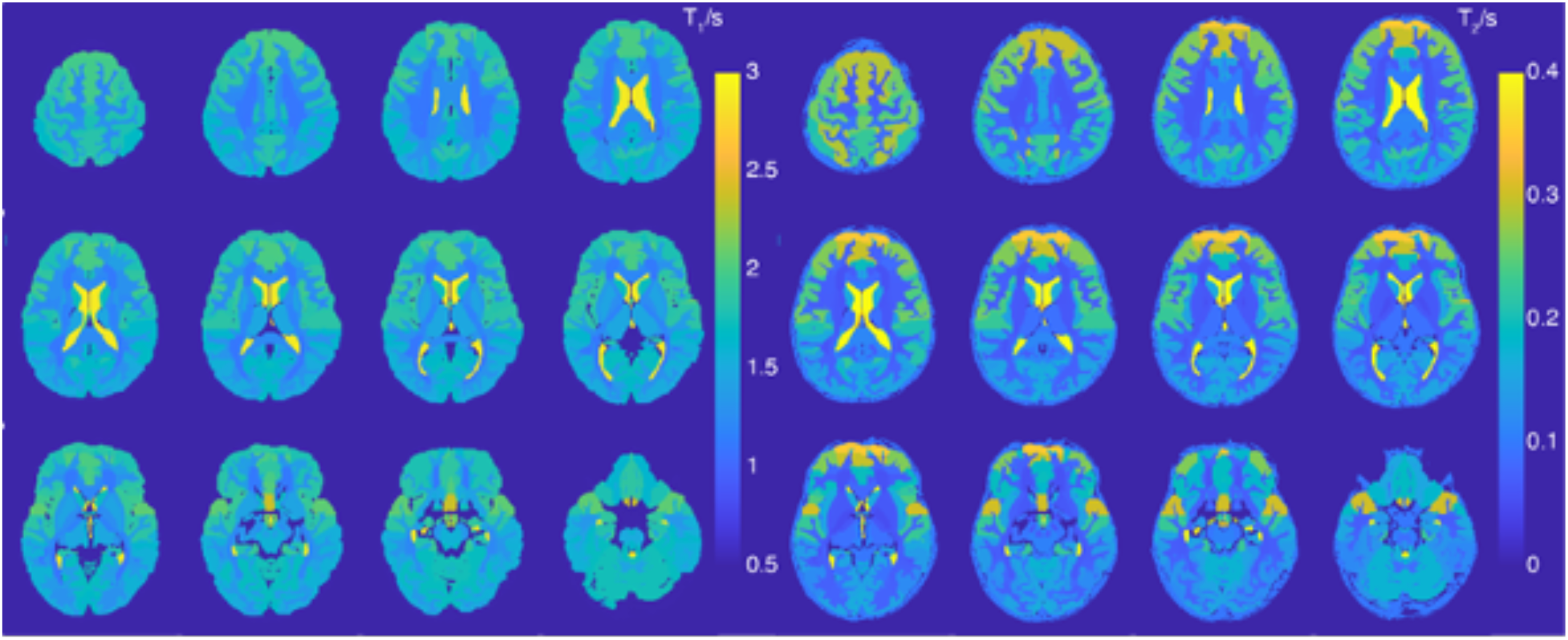
Age-30 atlas of predicted water *T*_1_ and *T*_2_ values for each anatomical parcel.

The standard Osprey workflow includes segmentation of the voxel with the Statistical Parametric Mapping (SPM12) (26) segmentation function. The ‘constant-value’ water-referenced relaxation corrections assumes literature values for water *T*_1_ and *T*_2_ for each segmented tissue-type *k* (where *k* is GM, WM or CSF). *T*_1w*k*_ and T_2w*k*_. Metabolite concentrations, *c*_met_, are then calculated from the linear-combination-model basis function amplitude, *S*_met_, according to:

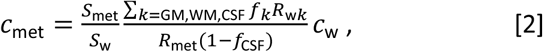

in which *S*_w_ is the amplitude of the water reference signal, *c*_w_ is the molal concentration of pure MR-visible water (55.5 mM), and the water relaxation factors *R*_w*k*_ combine *T*_1*wk*_ saturation and *T*_2*wk*_ decay terms:

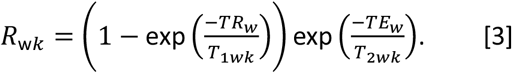

The *R*_met_ term has equivalent terms for metabolite relaxation, and the signal fractions, *f*_*k*_, for each tissue *k* are calculated from the volume fractions *f*_vol,*k*_:

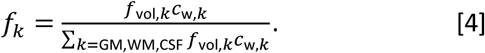

We have chosen to integrate the atlas into this workflow in a way that maintains continuity with this approach, i.e. using the atlas to extract location- and age-appropriate values for *T*_1w*k*_ and *T*_2w*k*_.

The voxel mask image generated by Osprey can then be resliced to the same image raster as the atlas_130.nii, so that every microvoxel within the voxel mask image can be assigned to a parcel label in the atlas image. Then, for each relevant parcel *j*, we calculate a predicted *T*_1j_ and *T*_2j_ based on the subject age and the model in Equation 1. We then average across parcels of a given tissue type, k, weighting by the number of micro-voxels *n*_*j*_ of each parcel within the voxel, e.g.

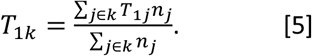

These predicted values for *T*_1w*k*_ and *T*_2w*k*_ can then be entered into Equation 2, replacing the standard literature values. The use of atlas-based water relaxation values in Osprey is enabled by a job-file flag, *‘MRSCont*.*opts*.*quantify*.*RelaxationAtlas’*, with age correction controlled by a second flag, ‘*MRSCont*.*opts*.*quantify*.*RelaxationAtlasAge’*. If the latter flag is enabled, participant age is drawn from a user-designated CSV file containing cohort characteristics, including age, to perform the age-adjusted corrections.

## 3. Results

DESPOT data were successfully acquired in 101 subjects. For four subjects, the LDDMM co-registration was repeated with less flexible parameters to address the misalignment of the corpus callosum. Atlas images populated with the age-30 predicted values of *T*_1_ and *T*_2_ for each anatomical parcel are shown in Figure 2. Both *T*_1_ and *T*_2_ images show the expected GM: WM contrast.

Table 1 shows the average *T*_1_ and *T*_2_ across parcels, weighted by parcel size, separated as GM, WM, and CSF. These values are compared to the prior reference values.

**Table 1.** GM, WM, and CSF parcel average *T*_1_ and *T*_2_ weighted by parcel size and comparison with literature. Osprey values for GM and WM were adapted from reference (3), while CSF *T*_1_ and *T*_2_ were sourced from (28) and (29), respectively.

| | Mean Atlas $T_1$<br>(s) | Prior Reference<br>$T_1$ (s) | | Mean Atlas<br>$T_2$<br>(ms) | Prior Reference<br>$T_2$ (ms) | |
| --- | --- | --- | --- | --- | --- | --- |
|  |  | Used in<br>Osprey (6) | DESPOT1<br>(27) |  | Used in<br>Osprey | DESPOT2-FM (19) |
| GM | 1.744 | 1.331 | ~1.6 | 178 | 110 | ~77.3 |
| WM | 1.399 | 0.832 | ~1.18 | 96 | 79.2 | ~50 |
| CSF | 2.888 | 3.817 |  | 566 | 503 |  |

The complete ‘*LookUpTable*.*mat’* is included as Supplemental Table 1.

Linear models of relaxation time against age were significant for *T*_1_ in 21 of 130 parcels, mostly in frontal, posterior, and inferior WM, and for 82 of 130 parcels for *T*_2_, particularly in GM, and frontal and limbic WM. The age-slope atlas is shown in Figure 3 and indicates the general lengthening of relaxation times with age. A notable exception is the shortening of T_1_ with age in sub-cortical areas. A correlation of *T*_1_ against *T*_2_ across parcels is shown in Figure 4, showing some separation of GM parcels, WM parcels, and CSF.

**Figure 3.**
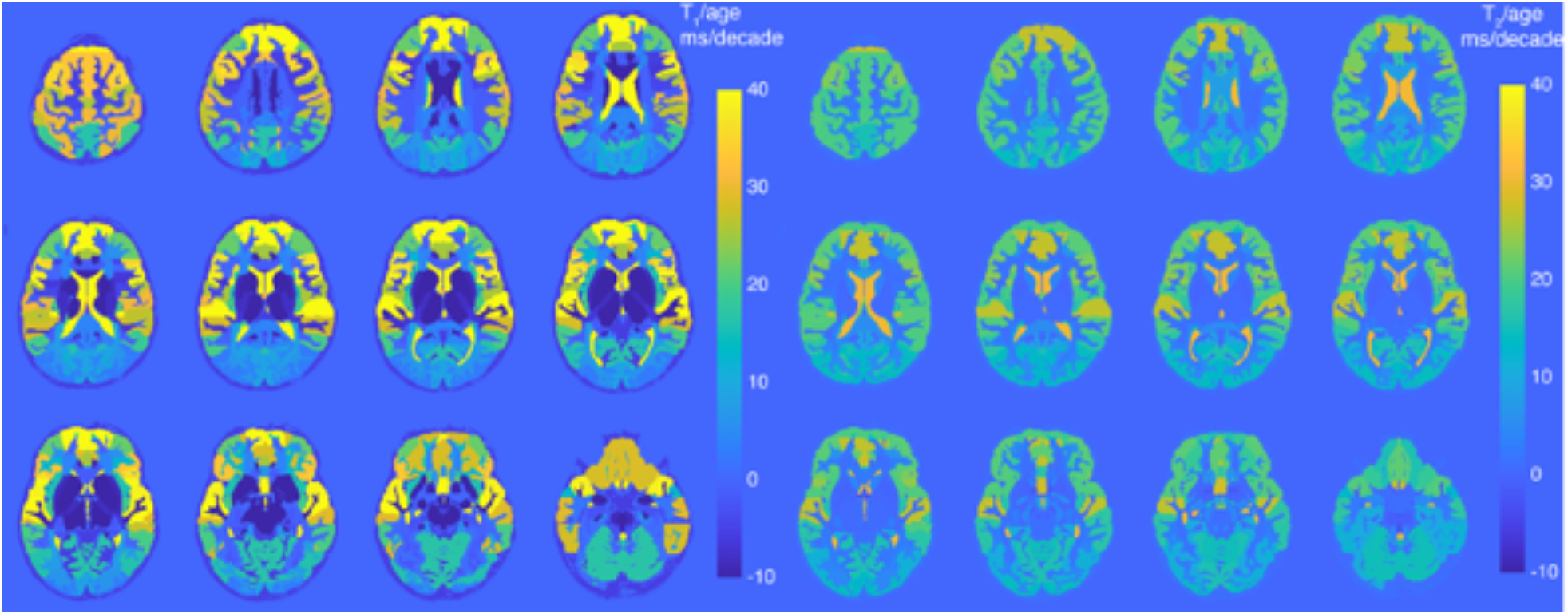
Age-slope atlases for predicted water *T*_1_ and *T*_2_ values for each anatomical parcel.

**Figure 4.**
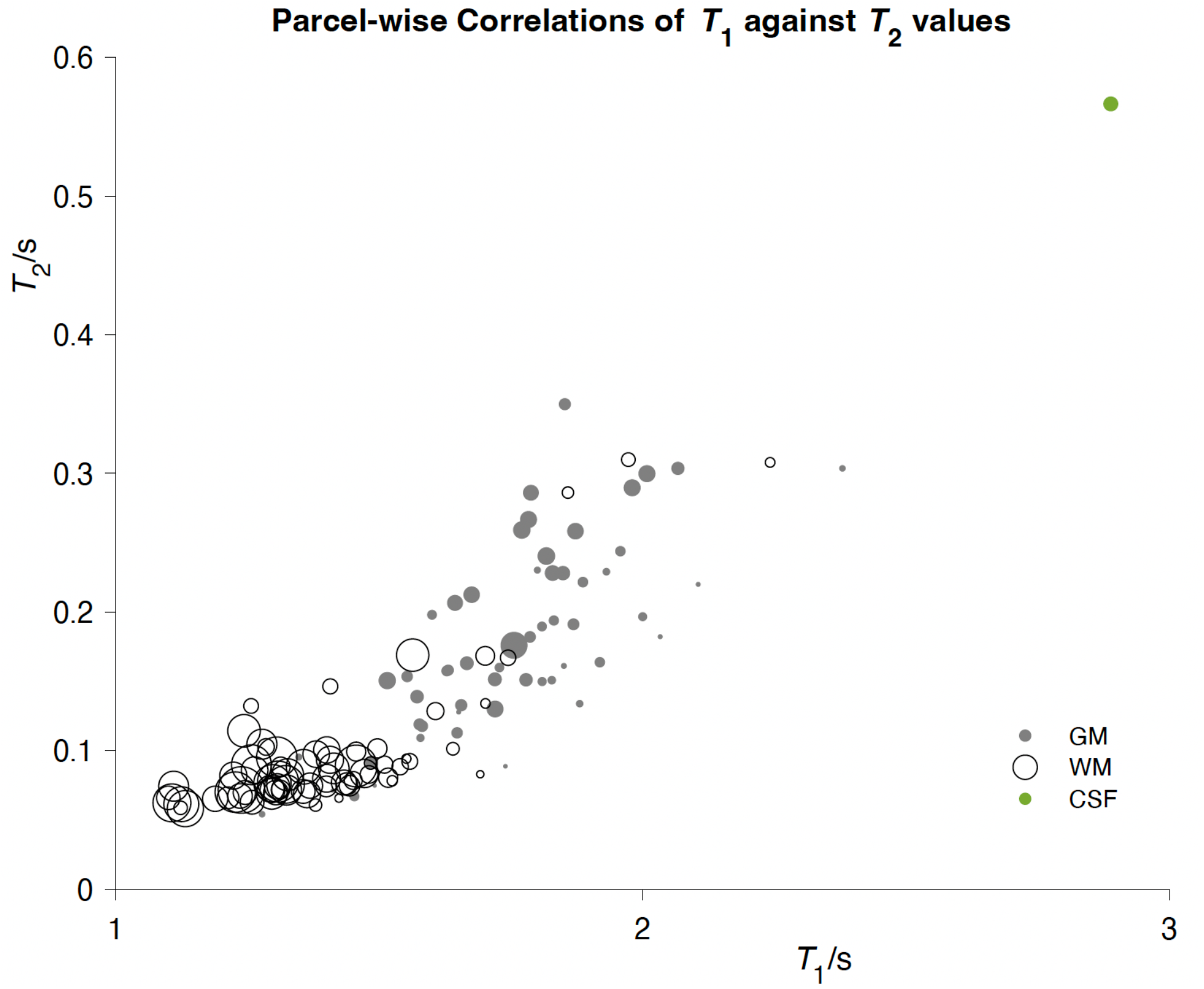
A parcel-wise correlation of *T*_1_ against *T*_2_ values colored as GM, WM and CSF. The point areas are scaled proportional to the parcel size.

## 4. Discussion

Water referencing in MRS is most often performed with corrections that use static literature references for relaxation times, despite the known age dependency of these parameters. This omission obfuscates true age effects on reported metabolite concentrations, which become entangled with the uncorrected differences in water relaxation. While the optimal solution would be to acquire subject-specific relaxometry data alongside MRS, this procedure is often impractically time-consuming, particularly in clinical settings. In this study, we have described a relaxometry dataset acquired in a structured age-span cohort to investigate the age dependency of water relaxation times. The dataset was co-registered to the JHU atlas by non-linear LDDMM deformation of each subject’s images, allowing the reference data to be summarized at the level of 130 anatomical parcels. We integrated these values into an automated workflow to facilitate age- and region-appropriate relaxation correction in MRS quantification.

The *T*_1_ values are in good agreement with literature values (27). For example, *T*_1_ values for the thalamus and occipital WM are reported as 1.5 s and 1.1 s, respectively, in the reference (27), compared to approximately 1.4 s and 1.3 s in the atlas. On average, the atlas reports GM and WM *T*_1_ values as 1.74 s and 1.39 s, while in (27) they were reported as 1.6 s and 1.18 s. On the other hand, the *T*_1_ values used in Osprey (3,6) are slightly lower, at 1.33 s for GM and 0.83 s for WM. Similarly, The *T*_2_ values show moderate agreement with prior literature. For instance, 3T DESPOT2 data (24) reports an average *T*_2_ of 50 ms in frontal WM across four subjects, while the atlas indicates an average of 60 ms for regions labeled as “Core Frontal WM.” The thalamus shows an average *T*_2_ of 73 ms compared to 91 ms in the atlas. Overall, the average *T*_2_ values in the atlas for GM and WM are 178 ms and 96 ms, respectively, whereas Osprey (3,6) uses 110 ms and 79.2 ms for these tissues.

Any attempt at water relaxometry must include the caveat that the values obtained depend strongly on the methodology applied (30–32). Here, we have used the DESPOT methodology which allows whole-brain relaxometry in under 11 minutes. Even more traditional methodologies, such as inversion recovery series for *T*_1_ and TE series for *T*_2_, will yield different values depending on the TIs and TE sampled, as will other relaxometry methods (including magnetization-prepared 2 rapid gradient echo (MP2RAGE), and multi-echo (ME) extension of MP2RAGE (MP2RAGEME) (33), QALAS (34,35), MR fingerprinting (36,37)). Ultimately, any attempt to describe the relaxation behavior of ∼10^19^ water molecules in a cubic millimeter of brain tissue in terms of just two parameters is an approximation. Future work might focus on acceleration to give better image resolution (ideally at 1 mm^3^ isotropic or better) and reduce the challenges of partial voluming.

While improved relaxometry resolution is desirable, the effective resolution of this atlas is probably more limited by co-registration than the resolution of the source images. The process of summarizing cohort data in an atlas inherently relies upon co-registering diverse brains. The question “Which point in my brain corresponds to this point in your brain?” is in some ways unanswerable, and within the limitations of MR-based co-registration, extremely challenging. In this work, we have used LDDMM, a non-linear co-registration that tries to maximize overlap to the EVE atlas brain. The LDDMM algorithm computes a diffeomorphic transformation between an atlas and a target image by solving a variational problem that aligns the images (23,24). This allows the encoding of unique shapes using vectors that are normal to the template’s outline. The algorithm optimizes this process via gradient descent, finding the optimal diffeomorphism that aligns the image to the atlas while minimizing a defined energy functional (i.e. smoothness of the deformation and the similarity), with adjustments for accuracy and robustness through cascading and multi-scale techniques. The proposed pipeline is imperfect due to anatomical diversity and mis-registration impacts the classification of CSF voxels as gray matter, which leads to biased *T*_1_ and particularly *T*_2_ values for the cortical GM parcels. With the current co-registration, it might be possible to use an additional masking step based on microvoxel segmentation classification at the single-subject level before averaging into parcels, as an additional filter to decrease bias.

This manuscript represents a significant step towards more accurate relaxation correction for MRS quantification. The underlying problem — that MR spectra are *T*_1_- and *T*_2_-weighted due to finite TR and TE — is one that MRS is stuck with. While water-referenced quantification of metabolites remains the consensus approach, the differential relaxation of water and metabolite signals must also be addressed. While a majority of work assumes literature values for water relaxation times, it has also been suggested to measure these parameters for each subject, either using multi-parametric and/or fingerprinting approaches (38–40), but it is likely that a cohort-reference-based approach will continue to be the best solution in terms of acquisition time and noise-bias trade-off. The approach used here for incorporating atlas values into the quantification equation was mainly used for the purposes of continuity with current practice. It is possible that, rather than averaging reference values within each tissue segment of the MRS voxel, a more refined approach would be to estimate the relaxation-weighted water signal within each microvoxel and average across these values.

## 5. Conclusion

The primary goal of this manuscript is to provide the means for future studies to quantify MRS data with region- and age-appropriate water reference values. Future work to improve upon the source dataset and the atlas-building process.

## Data availability

The data used in this study can be available from the corresponding author upon reasonable request.

## Acknowledgements

This work was supported by National Institutes of Health (NIH) grants R01 EB016089, R01 EB023963, R01 EB032788, R01 EB035529, R00 AG062230, R21 EB033516, K99 AG080084, K00 AG068440, P01AA029543, R01DK099334, and P41 EB031771.

## List of Abbreviations

bSSFP: balanced steady-state free precession
CSF: cerebrospinal fluid
DESPOT: driven-equilibrium single-pulse observazon of T1/T2
FA: flip angle
FOV: field of view
GM: gray ma|er
JHU: Johns Hopkins University
LDDMM: large deformazon diffeomorphic metric mapping
MNI-SS/EVE –: Montreal Neurological Insztute Single Subject Everything Parcellazon Map
MPRAGE: magnezzazon-prepared rapid gradient echo
MRS: magnezc resonance spectroscopy
Osprey: Open-source spectroscopy analysis so∼ware
SPGR: spoiled gradient-recalled echo
SPM: stazszcal parametric mapping
T_1_: longitudinal relaxazon zme
T_2_: transverse relaxazon zme
TE: echo zme
TR: repezzon zme
WM: white ma|er

## Notes

### Competing Interest Statement

The authors have declared no competing interest.

