## Supplementary Table 1 for "A Water Relaxation Atlas for Age- and Region-specific Metabolite Concentration Correction": Supplemental Table 1.docx

Supplemental Table 1: Look-up Table reproduced from Osprey.

| Index | Name | Class | Parcel size | *T*_1_ (s) | *T*_1_ slope (ms/year) | *T*_1__p | *T*_2_ (ms) | *T*_2_ slope (ms/year) | *T*_2__p | GM | WM |
| --- | --- | --- | --- | --- | --- | --- | --- | --- | --- | --- | --- |
| 1 | SFG | SFG | 46900 | 1.98 | 3.31 | 0.023 | 290 | 2.01 | 4.3x10^-8^ | 1 | 0 |
| 2 | SFG_PFC | SFG | 47313 | 2.008 | 5.19 | 0.003 | 300 | 2.7 | 7.63x10^-13^ | 1 | 0 |
| 3 | SFG_pole | SFG | 18228 | 1.852 | 4.69 | 0.03 | 350 | 2.01 | 2.90x^-5^ | 1 | 0 |
| 4 | MFG | MFG | 39281 | 1.873 | 3.51 | 0.013 | 258 | 2.3 | 4.06x10^-11^ | 1 | 0 |
| 5 | MFG_DPFC | MFG | 47533 | 1.783 | 2.19 | 0.203 | 267 | 2.1 | 1.29x10^-9^ | 1 | 0 |
| 6 | IFG_opercularis | IFG | 12563 | 1.958 | 3.97 | 0.006 | 244 | 2.42 | 3.55x10^-13^ | 1 | 0 |
| 7 | IFG_orbitalis | IFG | 12028 | 1.832 | 3.33 | 0.012 | 194 | 2.08 | 4.59x10^-13^ | 1 | 0 |
| 8 | IFG_triangularis IFG | IFG | 13135 | 1.887 | 3.70 | 0.005 | 222 | 2.3 | 9.16x10^-13^ | 1 | 0 |
| 9 | LFOG | OG | 16761 | 1.786 | 2.85 | 0.051 | 182 | 1.85 | 4.63x10^-11^ | 1 | 0 |
| 10 | MFOG | OG | 13826 | 1.628 | 2.88 | 0.1 | 158 | 1.58 | 5.12x10^-11^ | 1 | 0 |
| 11 | RG | RG | 17336 | 1.869 | 2.86 | 0.048 | 191 | 2.12 | 1.65x10^-13^ | 1 | 0 |
| 12 | PoCG | PoCG | 50484 | 1.771 | 3.06 | 0.017 | 259 | 2.01 | 4.08x10^-8^ | 1 | 0 |
| 13 | PrCG | PrCG | 51392 | 1.817 | 2.56 | 0.03 | 240 | 1.91 | 1.35x10^-9^ | 1 | 0 |
| 14 | SPG | SPG | 35253 | 1.788 | 3.02 | 0.057 | 286 | 1.99 | 8.02x10^-6^ | 1 | 0 |
| 15 | SMG | SMG | 35730 | 1.644 | 2.67 | 0.01 | 207 | 2.07 | 1.75x10^-11^ | 1 | 0 |
| 16 | AG | AG | 43840 | 1.676 | 1.88 | 0.109 | 213 | 2.02 | 3.11x10^-8^ | 1 | 0 |
| 17 | PrCu | PrCu | 34770 | 1.829 | 1.64 | 0.221 | 228 | 1.55 | 1.32x10^-4^ | 1 | 0 |
| 18 | STG | STG | 28177 | 1.849 | 4.84 | 2E-04 | 228 | 2.67 | 1.3x10^-14^ | 1 | 0 |
| 19 | STG_pole | STG | 22705 | 2.067 | 3.87 | 0.025 | 304 | 2.05 | 4.38x10^-7^ | 1 | 0 |
| 20 | MTG | MTG | 16199 | 1.553 | 2.92 | 0.006 | 154 | 1.75 | 3.84x10^-11^ | 1 | 0 |
| 21 | MTG_pole | MTG | 10675 | 1.809 | 1.28 | 0.421 | 190 | 1.31 | 3.57x10^-6^ | 1 | 0 |
| 22 | ITG | ITG | 18727 | 1.656 | -0.31 | 0.811 | 133 | 0.94 | 3.95x10^-7^ | 1 | 0 |
| 23 | PHG | Limbic | 5252 | 1.931 | 0.55 | 0.696 | 229 | 0.96 | 8.9x10^-4^ | 1 | 0 |
| 24 | ENT | Limbic | 2380 | 1.851 | 0.20 | 0.885 | 161 | 0.67 | 3.41x10^-2^ | 1 | 0 |
| 25 | FuG | Temporal | 40939 | 1.72 | 0.10 | 0.92 | 130 | 1.01 | 7.22x10^-7^ | 1 | 0 |
| 26 | SOG | Occipital | 10656 | 1.6 | 1.21 | 0.318 | 198 | 1.76 | 2.07x10^-6^ | 1 | 0 |
| 27 | MOG | Occipital | 49204 | 1.515 | 0.25 | 0.792 | 151 | 1.42 | 1.3x10^-7^ | 1 | 0 |
| 28 | IOG | Occipital | 17705 | 1.577 | 1.63 | 0.093 | 119 | 1.14 | 1.63x10^-6^ | 1 | 0 |
| 29 | Cu | Occipital | 25237 | 1.666 | 0.90 | 0.403 | 163 | 1.3 | 9.12x10^-7^ | 1 | 0 |
| 30 | LG | Occipital | 25930 | 1.72 | 1.66 | 0.147 | 151 | 1.57 | 1.09x10^-9^ | 1 | 0 |
| 31 | rostral_ACC | Cingulate | 8379 | 2 | 2.52 | 0.13 | 197 | 2.71 | 7.71X10^-12^ | 1 | 0 |
| 32 | subcallosal_ACC | Cingulate | 1302 | 2.033 | 0.61 | 0.73 | 182 | 1.67 | 3.75X10^-7^ | 1 | 0 |
| 33 | subgenual_ACC | Cingulate | 3259 | 2.379 | 5.76 | 0.011 | 304 | 2.79 | 2.06X10^-8^ | 1 | 0 |
| 34 | dorsal_ACC | Cingulate | 23654 | 1.779 | -0.08 | 0.946 | 151 | 1.76 | 8.63x10^-10^ | 1 | 0 |
| 35 | PCC | Cingulate | 15914 | 1.648 | 0.43 | 0.721 | 113 | 0.92 | 1.56x10^-7^ | 1 | 0 |
| 36 | Insula | Insula | 12563 | 1.919 | 1.58 | 0.219 | 164 | 2.02 | 2.87x10^-14^ | 1 | 0 |
| 37 | Amyg | Amyg | 4949 | 1.881 | -1.57 | 0.222 | 134 | 0.45 | 4.36x10^-3^ | 1 | 0 |
| 38 | Hippo | Hippo | 8718 | 1.809 | -0.26 | 0.833 | 150 | 1.02 | 1.03x10^-6^ | 1 | 0 |
| 39 | Caud | Caud | 9995 | 1.728 | -1.50 | 0.258 | 160 | 0.39 | 0.25 | 1 | 0 |
| 40 | Put | Put | 11362 | 1.453 | -1.50 | 0.088 | 67 | -0.004 | 0.53 | 1 | 0 |
| 41 | GP | GP | 3215 | 1.278 | -1.42 | 0.082 | 54 | 0.032 | 0.915 | 1 | 0 |
| 42 | Thalamus | Thalamus | 22283 | 1.486 | -1.30 | 0.205 | 91 | 0.071 | 0.496 | 1 | 0 |
| 43 | Hypo Thalamus | Basal Forebrain | 1223 | 2.105 | -1.48 | 0.527 | 220 | 1.03 | 0.038 | 1 | 0 |
| 44 | Mynert | Basal Forebrain | 674 | 1.492 | -0.99 | 0.43 | 75 | 0.047 | 0.765 | 1 | 0 |
| 45 | Nuc Accumbens | Basal Forebrain | 942 | 1.74 | -3.33 | 0.029 | 89 | -0.17 | 0.576 | 1 | 0 |
| 46 | RedNc | midbrain | 459 | 1.411 | -1.10 | 0.277 | 71 | -0.049 | 0.594 | 1 | 0 |
| 47 | Snigra | midbrain | 366 | 1.26 | -1.52 | 0.106 | 64 | -0.11 | 0.553 | 1 | 0 |
| 48 | Cerebellum | Cerebellum | 142477 | 1.756 | 1.68 | 0.129 | 176 | 1.48 | 5.18x10^-10^ | 1 | 0 |
| 49 | CP | midbrain | 4029 | 1.348 | -1.02 | 0.284 | 95 | 0.42 | 0.113 | 1 | 0 |
| 50 | Midbrain | midbrain | 6619 | 1.578 | -1.33 | 0.229 | 109 | 0.13 | 0.326 | 1 | 0 |
| 51 | CST | Pons | 3234 | 1.497 | -0.83 | 0.411 | 102 | -0.12 | 0.285 | 0 | 1 |
| 52 | SCP | Pons | 2966 | 1.701 | -0.28 | 0.83 | 168 | 0.43 | 0.053 | 0 | 1 |
| 53 | MCP | Pons | 14017 | 1.564 | -0.78 | 0.484 | 169 | 0.2 | 0.596 | 0 | 1 |
| 54 | PCT | Pons | 2206 | 1.51 | -1.14 | 0.259 | 90 | -0.0946 | 0.389 | 0 | 1 |
| 55 | ICP | Medulla | 2171 | 1.607 | -1.20 | 0.29 | 129 | 0.0566 | 0.9 | 0 | 1 |
| 56 | ML | Pons | 1682 | 1.558 | -0.57 | 0.604 | 92 | 0.1 | 0.79 | 0 | 1 |
| 57 | Pons | Pons | 1300 | 1.651 | -1.19 | 0.326 | 128 | -0.29 | 0.086 | 1 | 0 |
| 58 | Medulla | Medulla | 4105 | 1.8 | 0.55 | 0.696 | 230 | 0.13 | 0.913 | 1 | 0 |
| 59 | ACR | Core Frontal WM | 18515 | 1.132 | 0.39 | 0.565 | 58 | 0.11 | 0.195 | 0 | 1 |
| 60 | SCR | Core Frontal WM | 20745 | 1.107 | -0.04 | 0.948 | 62 | 0.12 | 0.021 | 0 | 1 |
| 61 | PCR | Core Posterior WM | 5832 | 1.1 | 0.24 | 0.735 | 66 | 0.11 | 0.376 | 0 | 1 |
| 62 | GCC | Corpus Callosum | 8750 | 1.223 | -0.12 | 0.9 | 82 | 0.27 | 0.117 | 0 | 1 |
| 63 | BCC | Corpus Callosum | 11469 | 1.278 | -1.38 | 0.362 | 105 | 0.81 | 0.031 | 0 | 1 |
| 64 | SCC | Corpus Callosum | 14211 | 1.244 | 0.39 | 0.723 | 114 | 0.53 | 0.047 | 0 | 1 |
| 65 | TAP | Inferior WM | 1295 | 1.257 | 1.01 | 0.527 | 132 | 0.92 | 0.063 | 0 | 1 |
| 66 | ALIC | Inferior WM | 6041 | 1.259 | -1.44 | 0.069 | 63 | 0.0282 | 0.969 | 0 | 1 |
| 67 | PLIC | Inferior WM | 7246 | 1.189 | -0.79 | 0.261 | 65 | 0.0644 | 0.915 | 0 | 1 |
| 68 | RLIC | Core Inferior WM | 4741 | 1.213 | -0.66 | 0.398 | 66 | 0.0836 | 0.122 | 0 | 1 |
| 69 | EC | Core Inferior WM | 8912 | 1.363 | -0.83 | 0.295 | 69 | 0.0824 | 0.717 | 0 | 1 |
| 70 | CGC | Limbic WM | 12080 | 1.296 | -1.69 | 0.069 | 69 | 0.0551 | 0.61 | 0 | 1 |
| 71 | CGH | Limbic WM | 3140 | 1.456 | -1.73 | 0.144 | 99 | 0.00951 | 0.89 | 0 | 1 |
| 72 | Fx/ST | Limbic WM | 3226 | 1.313 | -0.19 | 0.838 | 89 | 0.3 | 7.2x10^-3^ | 0 | 1 |
| 73 | Fx | Limbic WM | 983 | 1.973 | 3.03 | 0.252 | 310 | 2.2 | 3.2x10^-3^ | 0 | 1 |
| 74 | IFO | Core Inferior WM | 3981 | 1.444 | -1.20 | 0.211 | 75 | 0.0908 | 0.487 | 0 | 1 |
| 75 | PTR | Core Inferior WM | 11451 | 1.11 | 0.81 | 0.24 | 74 | 0.3 | 2.2x10^-3^ | 0 | 1 |
| 76 | SS | Core Inferior WM | 6668 | 1.236 | -0.05 | 0.948 | 67 | 0.17 | 0.015 | 0 | 1 |
| 77 | SFO | Core Inferior WM | 933 | 1.123 | -0.12 | 0.857 | 59 | 0.14 | 6.3x10^-3^ | 0 | 1 |
| 78 | SLF | Peripheral Parietal WM | 16976 | 1.125 | -0.13 | 0.854 | 61 | 0.0887 | 0.114 | 0 | 1 |
| 79 | UNC | Core Inferior WM | 715 | 1.64 | -0.59 | 0.586 | 101 | 0.1 | 2.5x10^-3^ | 0 | 1 |
| 80 | Ansa Lenticularis | Basal Forebrain | 685 | 1.484 | -1.22 | 0.255 | 91 | 0.55 | 3.1x10^-3^ | 0 | 1 |
| 81 | Anterior Com. | Basal Forebrain | 204 | 1.702 | -4.68 | 0.012 | 134 | -0.47 | 0.244 | 0 | 1 |
| 82 | Lenticular Fasc. | Basal Forebrain | 662 | 1.379 | -1.93 | 0.09 | 61 | 0.16 | 0.417 | 0 | 1 |
| 83 | Olfactory Radiation | Basal Forebrain | 95 | 1.692 | -2.80 | 0.082 | 83 | -0.15 | 0.761 | 0 | 1 |
| 84 | Mammillary | Basal Forebrain | 215 | 2.242 | -1.65 | 0.585 | 308 | 2.3 | 6.5x10^-3^ | 0 | 1 |
| 85 | Optic Tract | xxxx | 471 | 1.858 | -0.71 | 0.758 | 286 | 1.5 | 0.0104 | 0 | 1 |
| 86 | Ventricles | CSF | 30674 | 2.888 | 9.66 | 0.004 | 566 | 3.06 | 1.13x10^-5^ | 0 | 0 |
| 87 | PVWf | Frontal WM | 2003 | 1.285 | 1.15 | 0.223 | 103 | 1.05 | 2.4x10^-4^ | 0 | 1 |
| 88 | PVWp | Posterior WM | 1404 | 1.407 | 2.02 | 0.12 | 146 | 1.38 | 1.2x10^-4^ | 0 | 1 |
| 89 | SFWM | Peripheral Frontal WM | 24369 | 1.227 | -0.52 | 0.515 | 70 | 0.11 | 0.379 | 0 | 1 |
| 90 | SFWM_PFC | Peripheral Frontal WM | 20487 | 1.322 | -0.07 | 0.936 | 76 | 0.28 | 0.0192 | 0 | 1 |
| 91 | SFWM_pole | Peripheral Frontal WM | 1547 | 1.745 | 1.35 | 0.314 | 167 | 1.43 | 4.74x10^-5^ | 0 | 1 |
| 92 | MFWM | Peripheral Frontal WM | 20026 | 1.298 | -0.56 | 0.492 | 76 | 0.26 | 0.0171 | 0 | 1 |
| 93 | MFWM_DPFC | Peripheral Frontal WM | 12098 | 1.414 | 0.57 | 0.52 | 87 | 0.59 | 1.46x10^-6^ | 0 | 1 |
| 94 | IFWM_opercularis | Peripheral Frontal WM | 6948 | 1.298 | -0.63 | 0.469 | 72 | 0.13 | 0.378 | 0 | 1 |
| 95 | IFWM_orbitalis | Peripheral Frontal WM | 6652 | 1.304 | 1.23 | 0.123 | 70 | 0.3 | 1x10^-3^ | 0 | 1 |
| 96 | IFWM_triangularis | Peripheral Frontal WM | 8546 | 1.294 | 0.01 | 0.986 | 73 | 0.17 | 0.161 | 0 | 1 |
| 97 | LFOWM | Peripheral Frontal WM | 4102 | 1.4 | 1.21 | 0.25 | 74 | 0.34 | 0.0319 | 0 | 1 |
| 98 | MFOWM | Peripheral Frontal WM | 2573 | 1.451 | 1.65 | 0.129 | 78 | 0.43 | 6.03x10^-3^ | 0 | 1 |
| 99 | RGWM | Peripheral Frontal WM | 3221 | 1.517 | 0.65 | 0.555 | 81 | 0.64 | 1.13x10^-4^ | 0 | 1 |
| 100 | PoCWM | Peripheral Parietal WM | 24488 | 1.306 | -0.16 | 0.87 | 95 | 4.1 | 0.049 | 0 | 1 |
| 101 | PrCWM | Peripheral Frontal WM | 34140 | 1.239 | -0.44 | 0.571 | 72 | 0.21 | 0.086 | 0 | 1 |
| 102 | SPWM | Peripheral Parietal WM | 23575 | 1.258 | 0.06 | 0.94 | 90 | 0.39 | 0.0117 | 0 | 1 |
| 103 | SMWM | Peripheral Parietal WM | 13484 | 1.326 | -0.11 | 0.897 | 83 | 0.36 | 3.74x10^-3^ | 0 | 1 |
| 104 | AWM | Peripheral Parietal WM | 16564 | 1.357 | -0.17 | 0.839 | 88 | 0.45 | 2.4x10^-3^ | 0 | 1 |
| 105 | PrCuWM | Peripheral Parietal WM | 8270 | 1.381 | -0.13 | 0.882 | 97 | 0.41 | 0.045 | 0 | 1 |
| 106 | STWM | Peripheral Temporal WM | 9394 | 1.4 | -1.14 | 0.262 | 80 | 0.15 | 0.283 | 0 | 1 |
| 107 | STWM_pole | Peripheral Temporal WM | 1882 | 1.54 | -1.46 | 0.216 | 88 | -0.11 | 0.492 | 0 | 1 |
| 108 | MTWM | Peripheral Temporal WM | 6076 | 1.356 | 0.21 | 0.813 | 71 | 0.27 | 0.023 | 0 | 1 |
| 109 | MTWM_pole | Peripheral Temporal WM | 2448 | 1.481 | -0.23 | 0.825 | 83 | 0.0881 | 0.377 | 0 | 1 |
| 110 | ITWM | Peripheral Temporal WM | 7017 | 1.433 | 0.23 | 0.802 | 77 | 0.23 | 0.0139 | 0 | 1 |
| 111 | FuWM | Peripheral Temporal WM | 9871 | 1.472 | -0.21 | 0.813 | 84 | 0.24 | 0.019 | 0 | 1 |
| 112 | SOWM | Peripheral Occipital WM | 8372 | 1.264 | 0.76 | 0.307 | 86 | 0.53 | 6.01x10^-4^ | 0 | 1 |
| 113 | MOWM | Peripheral Occipital WM | 25165 | 1.308 | 0.65 | 0.359 | 78 | 0.44 | 1.21x10^-5^ | 0 | 1 |
| 114 | IOWM | Peripheral Occipital WM | 7178 | 1.369 | 0.75 | 0.334 | 75 | 0.38 | 3.3x10^-4^ | 0 | 1 |
| 115 | CuWM | Peripheral Occipital WM | 8087 | 1.401 | 0.41 | 0.625 | 101 | 0.51 | 8.75x10^-4^ | 0 | 1 |
| 116 | LWM | Peripheral Occipital WM | 8757 | 1.407 | -0.53 | 0.567 | 93 | 0.24 | 0.118 | 0 | 1 |
| 117 | Rostral WM_ACC | Peripheral Limbic WM | 294 | 1.525 | -1.31 | 0.257 | 78 | 0.26 | 0.039 | 0 | 1 |
| 118 | Subcallosal WM_ACC | Peripheral Limbic WM | 110 | 1.424 | -0.21 | 0.854 | 66 | 0.0521 | 0.554 | 0 | 1 |
| 119 | Subgenualn WM_ACC | Peripheral Limbic WM | 170 | 1.552 | -0.23 | 0.884 | 94 | 0.31 | 0.091 | 0 | 1 |
| 120 | Dorsal WM_ACC | Peripheral Limbic WM | 2806 | 1.313 | -1.37 | 0.112 | 72 | 0.16 | 0.163 | 0 | 1 |
| 121 | PCCWM | Peripheral Limbic WM | 6165 | 1.246 | -0.27 | 0.752 | 70 | 0.12 | 0.157 | 0 | 1 |
| 122 | Cerebellum WM | Cerebellum | 26287 | 1.456 | 0.00 | 0.998 | 89 | 0.18 | 0.068 | 0 | 1 |
| 123 | CSF | CSF | 424818 | 0.517 | -0.56 | 0.208 | 97 | 0.12 | 0.109 | 0 | 0 |
| 124 | Pins | Insula | 7265 | 1.828 | 1.96 | 0.127 | 151 | 1.64 | 1.22x10^-13^ | 1 | 0 |
| 125 | PSTG | STG | 14920 | 1.631 | 2.85 | 0.004 | 158 | 2.06 | 1.71x10^-14^ | 1 | 0 |
| 126 | PMTG | MTG | 24166 | 1.572 | 2.00 | 0.03 | 139 | 1.69 | 3.03x10^-11^ | 1 | 0 |
| 127 | PITG | ITG | 17388 | 1.582 | 2.81 | 0.015 | 118 | 1.16 | 7.89x10^-9^ | 1 | 0 |
| 128 | PSTGWM | Peripheral Temporal WM | 7197 | 1.306 | -0.71 | 0.422 | 74 | 0.16 | 0.181 | 0 | 1 |
| 129 | PMTGWM | Peripheral Temporal WM | 11648 | 1.324 | 0.02 | 0.981 | 72 | 0.29 | 4.99x10^-3^ | 0 | 1 |
| 130 | PITGWM | Peripheral Temporal WM | 4781 | 1.438 | 0.79 | 0.346 | 76 | 0.37 | 6.03x10^-4^ | 0 | 1 |

List of Abbreviations

- ACR - Anterior Commissure
- ALIC - Anterior Limb of Internal Capsule
- Amyg - Amygdala
- Anterior Com - Anterior Commissure
- AWM - Angular White Matter
- Caud - Caudate Nucleus
- CGC - Cingulum Gyrus
- CGH - Cingulate Gyrus
- Cu - Cuneus
- CuWM - Cuneus White Matter
- CSF - Cerebrospinal Fluid
- dorsal_ACC - Dorsal Anterior Cingulate Cortex
- dorsal WM_ACC - Dorsal White Matter Anterior Cingulate Cortex
- EC - Entorhinal Cortex
- Fx - Fornix
- Fx/ST - Fornix/Splenium of Corpus Callosum
- FuG - Fusiform Gyrus
- FuWM - Fusiform White Matter
- GP - Globus Pallidus
- GCC - Genu of Corpus Callosum
- IFG_opercularis - Inferior Frontal Gyrus, Opercularis
- IFG_orbitalis - Inferior Frontal Gyrus, Orbitalis
- IFG_triangularis - Inferior Frontal Gyrus, Triangularis
- IFWM_opercularis - Inferior Frontal White Matter, Opercularis
- IFWM_orbitalis - Inferior Frontal White Matter, Orbitalis
- IFWM_triangularis - Inferior Frontal White Matter, Triangularis
- Insula - Insular Cortex
- IOG - Inferior Occipital Gyrus
- IOWM - Inferior Occipital White Matter
- ITG - Inferior Temporal Gyrus
- ITWM - Inferior Temporal White Matter
- LFOG - Lateral Fronto-Occipital Gyrus
- LFOWM - Lateral Fronto-Occipital White Matter
- LG - Lingual Gyrus
- LWM - Lingual White Matter
- Mammillary - Mammillary Bodies
- MFOWM - Medial Fronto-Occipital White Matter
- MFWM - Middle Frontal White Matter
- MFWM_DPFC - Middle Frontal White Matter, Dorsal Prefrontal Cortex
- MFOG - Medial Fronto-Occipital Gyrus
- MFG - Middle Frontal Gyrus
- MFG_DPFC - Middle Frontal Gyrus, Dorsal Prefrontal Cortex
- MOG - Middle Occipital Gyrus
- MTG - Middle Temporal Gyrus
- MTG_pole - Middle Temporal Gyrus, Pole
- MTWM - Middle Temporal White Matter
- MTWM_pole - Middle Temporal White Matter, Pole
- Mynert - Mynert Nucleus (Basal Forebrain)
- NucAccumbens - Nucleus Accumbens
- PCC - Posterior Cingulate Cortex
- PCCWM - Posterior Cingulate Cortex White Matter
- PHG - Parahippocampal Gyrus
- PrCG - Precentral Gyrus
- PrCu - Precuneus
- PrCuWM - Precuneus White Matter
- Put - Putamen
- RedNc - Red Nucleus
- RG - Recurrent Gyrus
- RGWM - Recurrent Gyrus White Matter
- rostral_ACC - Rostral Anterior Cingulate Cortex
- rostralWM_ACC - Rostral White Matter Anterior Cingulate Cortex
- SFO - Superior Fronto-Occipital Fasciculus
- SFG - Superior Frontal Gyrus
- SFG_PFC - Superior Frontal Gyrus, Prefrontal Cortex
- SFG_pole - Superior Frontal Gyrus, Pole
- SFWM - Superior Frontal White Matter
- SFWM_PFC - Superior Frontal White Matter, Prefrontal Cortex
- SFWM_pole - Superior Frontal White Matter, Pole
- SMG - Supramarginal Gyrus
- SMWM - Supramarginal White Matter
- Snigra - Substantia Nigra
- SOG - Superior Occipital Gyrus
- SLF - Superior Longitudinal Fasciculus
- SOWM - Superior Occipital White Matter
- SPG - Superior Parietal Gyrus
- SPWM - Superior Parietal White Matter
- STG - Superior Temporal Gyrus
- STG_pole - Superior Temporal Gyrus, Pole
- STWM - Superior Temporal White Matter
- STWM_pole - Superior Temporal White Matter, Pole
- TAP - Thalamic Anterior Pole
- PTR - Posterior Thalamic Radiation
- PVWf - Posterior Ventral White Matter Fasciculus
- PVWp - Posterior Ventral White Matter Peduncle
- PSTG - Posterior Superior Temporal Gyrus
- PSTGWM - Posterior Superior Temporal Gyrus White Matter
- PMTG - Posterior Middle Temporal Gyrus
- PMTGWM - Posterior Middle Temporal Gyrus White Matter
- PITG - Posterior Inferior Temporal Gyrus
- PITGWM - Posterior Inferior Temporal Gyrus White Matter
- Subcallosal_ACC - Subcallosal Anterior Cingulate Cortex
- Subgenual_ACC - Subgenual Anterior Cingulate Cortex
- subcallosal WM_ACC - Subcallosal White Matter Anterior Cingulate Cortex
- Subgenual WM_ACC - Subgenual White Matter Anterior Cingulate Cortex
- SCC - Splenium of Corpus Callosum
- SCR - Superior Commissure
- SFO - Superior Fronto-Occipital Fasciculus
- SLF - Superior Longitudinal Fasciculus
- UNC - Uncinate Fasciculus
- Lenticular Fasc - Lenticular Fasciculus
- SFWM - Superior Frontal White Matter
- MFWM - Middle Frontal White Matter
- MFWM_DPFC - Middle Frontal White Matter, Dorsal Prefrontal Cortex
- IFWM_opercularis - Inferior Frontal White Matter, Opercularis
- IFWM_orbitalis - Inferior Frontal White Matter, Orbitalis
- IFWM_triangularis - Inferior Frontal White Matter, Triangularis
- LFOWM - Lateral Fronto-Occipital White Matter
- MFOWM - Medial Fronto-Occipital White Matter
- RGWM - Recurrent Gyrus White Matter
- PoCWM - Postcentral Cortex White Matter
- PrCWM - Precentral Cortex White Matter
- SPWM - Superior Parietal White Matter
- SMWM - Supramarginal White Matter
- AWM - Angular White Matter
- PrCuWM - Precuneus White Matter
- STWM - Superior Temporal White Matter
- STWM_pole - Superior Temporal White Matter, Pole
- MTWM - Middle Temporal White Matter
- MTWM_pole - Middle Temporal White Matter, Pole
- ITWM - Inferior Temporal White Matter
- FuWM - Fusiform White Matter
- SOWM - Superior Occipital White Matter
- MOWM - Middle Occipital White Matter
- IOWM - Inferior Occipital White Matter
- LWM - Lingual White Matter
- Rostral WM_ACC - Rostral White Matter Anterior Cingulate Cortex
- Subcallosal WM_ACC - Subcallosal White Matter Anterior Cingulate Cortex
- Subgenual WM_ACC - Subgenual White Matter Anterior Cingulate Cortex
- Dorsal WM_ACC - Dorsal White Matter Anterior Cingulate Cortex
- PCCWM - Posterior Cingulate Cortex White Matter
